# Grandmaternal smoking during pregnancy is associated with differential DNA methylation in their grandchildren

**DOI:** 10.1101/2021.04.22.440892

**Authors:** Sarah Holmes Watkins, Yasmin Iles-Caven, Marcus Pembrey, Jean Golding, Matthew Suderman

**Affiliations:** University of Bristol, MRC Integrative Epidemiology Unit, Population Health Sciences, Bristol Medical School, Bristol, United Kingdom; University of Bristol, Centre for Academic Child Health, Population Health Sciences, Bristol Medical School, Bristol, United Kingdom

**Author notes:** Corresponding author: Sarah Watkins. **Funding** The UK Medical Research Council and Wellcome (Grant ref: 217065/Z/19/Z) and the University of Bristol provide core support for ALSPAC. This research was made possible through the support of a grant from the John Templeton Foundation (60828).

**Keywords:** ALSPAC, grandmaternal smoking, DNA methylation, transgenerational effects, prenatal smoking

## Abstract

The idea that information can be transmitted to subsequent generation(s) by epigenetic means has been studied for decades but remains controversial in humans. Epidemiological studies have established that grandparental exposures are associated with health outcomes in their grandchildren, often with sex-specific effects; however the mechanism of transmission is still unclear. We conducted Epigenome Wide Association Studies (EWAS) to test whether grandmaternal smoking during pregnancy is associated with altered DNA methylation (DNAm) in their adolescent grandchildren. We used data from a birth cohort, with discovery and replication datasets of 1225 and 708 individuals (respectively), aged 15-17 years, and tested replication in the same individuals at birth and 7 years. We show for the first time that DNAm at a small number of loci is associated with grandmaternal smoking in humans, and their locations in the genome suggest hypotheses of transmission. We observe and replicate sex-specific associations at two sites on the X chromosome, one located in an imprinting control region and both within transcription factor binding sites (TFBSs). In fact, we observe enrichment for TFBSs among the CpG sites with the strongest associations, suggesting that TFBSs may be a mechanism by which grandmaternal exposures influence offspring DNA methylation. There is limited evidence that these associations appear at earlier timepoints, so effects are not static throughout development. The implication of this work is that effects of smoking during pregnancy may induce DNAm changes in later generations and that these changes are often sex-specific, in line with observational associations.

## Introduction

The idea that information can be transmitted to subsequent generation(s) by epigenetic means remains controversial in humans (1). The terminology used in the literature on this topic is not always consistent; here we use the term transgenerational to include all transmissions from one generation to subsequent generations. Of all epigenetic mechanisms that might be involved in transmission of information between generations, DNA methylation (DNAm) is a strong candidate because it is heritable over cell division. A frequent argument against this is the two widespread phases of global de-methylation followed by re-methylation that all humans undergo in germ cells, and then immediately post-fertilization at the blastocyst stage (which is necessary to allow cells to become pluripotent (2, 3)). However it has been shown that in human germ cells some genomic regions escape de-methylation (4), and imprinting control regions (ICRs) are not subject to de-methylation or re-methylation in the early embryo (5). Reports are also emerging of mechanisms which may preserve or restore DNAm at certain loci during the phase of germ cell de-methylation - such as transcription factors (TFs) (6, 7).

Most work on transgenerational responses in humans stems from epidemiological studies which report associations between grandparental exposures or experiences and grandchild health outcomes. Effects are often sex specific and unique to either the maternal or paternal line (8, 9). Tobacco smoke exposure accounts for a significant proportion of transgenerational studies in humans; this is because in human cohort studies records of smoking behaviour are commonly available, and compared to other exposures and lifestyle factors smoking behaviour is relatively easy to objectively and accurately recall and record, even by family members.

Grandmaternal smoking during pregnancy has been associated with a number of health outcomes in their grandchildren: paternal grandmother’s smoking during pregnancy is associated with greater fat mass in their adult granddaughters, but not grandsons (10), and with reduced prevalence of myopia in their grandchildren up to 7 years of age (with stronger effects in the grandsons); this study also demonstrated that early-onset myopia is associated with DNAm at multiple loci (11). Maternal grandmother smoking during pregnancy is associated with higher birth weight, and subsequent greater lean mass and higher cardiovascular fitness, in their grandsons (12, 13). It has been well established that smoking is associated with differences in DNAm, both in the individual (14, 15) and in the offspring of mothers who smoke during pregnancy (16, 17); currently one published study has assessed the association of grandmaternal smoking with DNAm in their grandchildren, at 26 DNAm sites that were known to be associated with prenatal smoke exposure. However none of these sites were found to be associated with grandmaternal smoking (18).

Here we test the hypothesis that grandmaternal smoking during pregnancy is associated with differences in DNAm in their grandchildren, at over 450,000 sites across the genome. We utilise the Avon Longitudinal Study of Parents and Children (ALSPAC) cohort (19), which has both DNAm data and detailed information about ancestral smoking. Almost 3,000 methylomes have now been assayed for the index children at 15 years of age, making this the largest human cohort available to assess transgenerational epigenetic inheritance.

## Methods

### Cohort description

We used two DNAm datasets from the ALSPAC cohort; please see supplementary methods for a detailed cohort description. Our discovery data were a newly generated dataset of 1869 individuals at 15-17 years of age, who had their methylomes assayed on the Illumina EPIC Human Methylation microarray (EPIC array). Replication analyses utilised the original subsample of ALSPAC with DNAm data (known as the Accessible Resource for Integrated Epigenomic Studies, ARIES), assayed on the Illumina 450K Human Methylation microarray (450k array) (20). Although replication is ideally conducted in a separate dataset, we could identify no other available DNAm datasets of a similar age with ancestral smoking data.

### Ancestral smoking data

We determined whether the maternal and paternal grandmothers of the ALSPAC study children smoked during pregnancy using questionnaires completed by the study mother and father. Maternal and paternal lines were tested separately. The variable we used was a categorical ‘Yes’ or ‘No’; see supplementary methods for details on how this was created.

### Study exclusions

To attempt to detect only grandmaternal effects, we excluded all individuals whose mother reported smoking whilst pregnant, and all adolescents who reported smoking themselves. We also excluded a small number of individuals from the EPIC dataset who were of non-white ethnicity, as reduced rates of both maternal and paternal grandmother smoking were associated with non-white ethnicity; please see supplementary methods for details.

### New ALSPAC methylomes assayed on EPIC array

DNA methylation profiles were generated as previously described but using the Illumina Infinium MethylationEPIC Beadchip (EPIC array) rather than then the Illumina Infinium HumanMethylation450k Beadchip (450k array) (21). Briefly, following DNA extraction, DNA was bisulfite converted using the Zymo EZ DNA Methylation™ kit (Zymo, Irvine, CA), and DNA methylation was measured using EPIC arrays. Arrays were scanned using Illumina iScan, and the initial quality review was carried out using GenomeStudio. A wide range of batch variables were recorded in a purpose-built laboratory information management system (LIMS). Additional quality control and normalization was carried out using the *meffil* R package. Of 1885 initial samples, 16 were found to have an unacceptably high proportion of undetected probes (proportion > 10% with detection p-value < 0.01). The remaining 1869 were normalized using functional normalization (22) as implemented in *meffil* with quantiles adjusted using 20 control probe principal components and slide as a random effect.

### Original ARIES DNAm data assayed on 450k array

921 samples were available for the 15–17-year-olds from the original ARIES DNAm dataset, along with 849 individuals at birth, and 910 individuals at 7 years. Consent for biological samples was collected in accordance with the Human Tissue Act (2004). Processing, extraction, and quality control of DNAm data has been described in detail for these samples (20), as have normalisation and outlier removal procedures (21). 21 further individuals were removed as they were the only sample on a slide, preventing their adjustment for slide effects.

### Filtering DNAm sites

All sites on the X and Y chromosomes were removed from the analyses using all individuals; the Y chromosome was removed from the single sex analyses. No other sites were removed, but results were checked against probes flagged as being cross-reactive or having a SNP at the CpG site, in the single base extension, or in the probe body (23). Results were also checked for being located in the highly polymorphic human leukocyte antigen (HLA) region.

### EWAS

Six EWAS were performed in both the discovery (EPIC) and replication (450k) datasets, testing the association of DNAm with maternal grandmother smoking in all individuals, in females, and in males, and with paternal grandmother smoking, in all individuals, in females, and in males. EWAS were conducted using the R package meffil (21). A genome-wide significance threshold of p<9e-08 was used for the EPIC analyses (24), and p<2.4e-07 for the 450k (25). Covariates for all EWAS were: age at DNAm sample; sex (for the analyses with both sexes); batch effects (plate for the EPIC samples, slide for the 450k samples); and cell count proportions estimated using a deconvolution algorithm (26) implemented in meffil, based on the “blood gse35069 complete” cell type reference. Because known covariates can be imperfect and miss sources of unwanted variation, we conducted a sensitivity analysis adjusting for surrogate variables (using surrogate variable analysis (SVA)(27) as implemented in meffil) where we assessed correlation between effect sizes of the SVA and known covariates models. As there was high correlation between effect sizes (>0.97) the known covariates model was used for all analyses – the high correlation suggests that the main model accounted for all substantial sources of DNAm variation, and SVA risks removing biologically interesting sources of variation in the data.

### Testing for replication

We used three complementary approaches to test for replication of the sites most strongly associated with the exposure in the discovery dataset, as no single measure can capture this. Firstly, we took the 25 top associated sites from the EPIC analyses that were also present on the 450k array and assessed them for association in the 450k analyses at the equivalent of p<0.05/25. Secondly, we correlated effect sizes between the discovery and replication datasets, for the top 10, 25, 50, 100 and 200 sites identified in each discovery EWAS. Finally, we conducted a binomial test for each discovery EWAS to ascertain whether the top 10, 25, 50, 100 and 200 sites replicated at p<0.05 with the same direction of effect.

### Meta-analysis

For each of the six EWAS (maternal grandmother smoking: all individuals, males, and females; and paternal grandmother smoking: all individuals, males, and females), we meta-analysed results from the EPIC and 450k analyses at 15-17 years, using all sites common to both arrays. We performed meta-analysis of the effect sizes and standard errors using METAL (28).

### X chromosome analysis

As previous work has identified that associations between ancestral exposures and health outcomes in later generations are often sex-specific, and X-inactivation is specific to females, we tested the hypothesis that sites on the X chromosome would associate with grandmaternal smoking during pregnancy. We tested the X chromosome separately for each sex-stratified EWAS and meta-analysis, adjusting for X chromosome significance (p<2.7e-06 in EPIC, p<4.5e-06 in 450k and p<4.9e-06 in the meta-analysis).

### Escapees analysis

As some DNAm sites have been shown to escape the wave of de-methylation in germ cells (4), we tested the hypothesis that these sites are associated with grandmaternal smoking during pregnancy. To do this we took the 116,618 regions of the genome that have been identified as escaping de-methylation (4) (which have recently been made available as supplementary material in a bioRxiv paper (29)). We identified all sites on the EPIC array that were within those genomic regions (n=36,051) and tested them for association with grandmaternal smoking at Bonferroni corrected significance (p<0.05/36051=1.4e-6).

### Imprinting control region analysis

As ICRs are not subject to the phases of de-methylation and re-methylation in the early embryo, we sought to test whether DNAm sites in identified regions might associate with grandmaternal smoking. We took the set of 984 DNAm sites present on the EPIC array identified as being within ICRs at FDR<0.05 (30). We tested these sites for association with grandmaternal smoking at Bonferroni corrected significance (p<0.05/984=5.1e-5). There were 29 ICR sites that overlapped with the escapees.

### Testing replication earlier in life

To ascertain whether any sites associated with grandmaternal smoking at 15-17 years are differentially methylated from birth, we repeated each EWAS using DNAm profiles for ALSPAC participants from blood samples collected at birth and 7 years (see supplementary table 1 for participant numbers). We included the same covariates as for the adolescents, aside from at birth where gestational age was substituted for age. As the birth and 7 years DNAm profiles were assayed from different sample types (blood spots and white cells at birth; white cells and whole blood at 7 years), sample type was also included as a covariate. In addition to using this analysis to assess replication of associations in the 15-17-year-olds, we assessed the opposite, replication of associations at the birth and age 7 in the 15-17-year-old discovery dataset.

**Table 1:** Summary of variables used in the analysis for participants with DNAm data assayed on the EPIC and 450k arrays. MGM = maternal grandmother, PGM = paternal grandmother. * percentages in this cell are calculated after omitting participants with missing ancestral smoking data.

| Dataset | Measure | MGM analysis | PGM analysis | Full cohort |
| --- | --- | --- | --- | --- |
| EPIC | N participants | 1225 | 1021 | 1869 |
|  | Grandmaternal smoking (% yes) | 30.9% | 38.4% | 34% MGM; 39% PGM* |
|  | Sex (% Female) | 50.7% | 51.6% | 52.4% |
|  | Age (mean(SD)) | 17.8 (0.4) | 17.8 (0.4) | 17.8 (0.5) |
| 450k | N participants | 708 | 601 | 910 |
|  | Grandmaternal smoking (% yes) | 29.9% | 36.6% | 31.7% MGM; 39.4% PGM* |
|  | Sex (% Female) | 51.1% | 51.6% | 51.6% |
|  | Age (mean(SD)) | 17.1 (1) | 17.1 (1) | 17.1 (1) |

**Table 2:** Sites which were found to associate with grandmaternal smoking in all analyses. Models were adjusted for all covariates (age/gestational age, predicted cell counts, and batch effects).

| Age (subset) | Array | Analysis | CpG | chr | position | p value | Effect size |
| --- | --- | --- | --- | --- | --- | --- | --- |
| 15-17 (females) | EPIC + 450k | Replication | cg19782749 | Xchr | 132091720 | 5.2e-05 (EPIC); 0.001 (450k) | 0.002 (EPIC); 0.001 (450k) |
| 15-17 (males) | EPIC | PGM (X chr) | cg27456137 | chrX | 129403024 | 1.9e-06 | -0.01 |
| 15-17 (all individuals) | EPIC | PGM (ICR) | cg15068552 | 7 | 130130203 | 2.2e-05 | -0.02 |
| Birth (all individuals) | 450k | MGM | cg19426678 | 12 | 117537404 | 2.1e-07 | -0.002 |
| Birth (females) | 450k | PGM | cg22682200 | 10 | 120001266 | 2.3e-09 | -0.05 |
| Birth (females) | 450k | PGM | cg26827966 | 5 | 172189374 | 5.9e-08 | -0.03 |

### Transcription factor binding site (TFBS) enrichment analysis

To test the hypothesis that differential DNAm associated with grandmaternal smoking might be mediated by TFs preserving or maintaining methylation status, we tested whether DNAm sites were located near TFBS more than expected by chance. To do this we took the top 25 sites from each discovery EWAS and tested them for TFBS enrichment against all sites on the EPIC array used in our EWAS (n sites=838,019) using LOLA locus overlap (31). We used the Encode TFBS (32, 33) region set created by the LOLA team, comprising ChIP-seq data on 161 TFs, which is available through http://lolaweb.databio.org. We tested 100bp on either side of the DNAm site, removing overlapping sites to prevent inflation of results. Results were reduced to TFBS measured in blood which were associated in at least one EWAS at p<0.05. To assess whether individual sites identified in the main analysis were associated with a TFBS, we used the hg19 version of the UCSC genome browser (34); https://genome-euro.ucsc.edu/.

### Enrichment of prenatal- and own smoking- associated sites

We tested the hypothesis that DNAm sites that are established as being associated with prenatal smoking and own smoking would be enriched in our EWAS associations, to ascertain whether transgenerational transmission might be related to these sites. To do this we evaluated statistical inflation of EWAS associations among the 568 DNAm sites (of which 540 were available on the EPIC array) previously reported to be associated with maternal prenatal smoking in cord blood (17), and the 2623 sites (2445 available on the EPIC array) reported to be associated with own smoking (35). For each, inflation beyond expected levels was evaluated by generating QQ plots and lambda values. We then used a one-sided Wilcoxon rank sum test to ask if DNAm sites associated with prenatal- and own-smoking had lower p-values in our EWAS than expected from a random selection.

### Enrichment of lean mass-associated sites

We finally sought to identify whether DNAm sites associated with grandmaternal smoking might be related to lean mass (a previously reported epidemiological association (13)). Although no published EWAS of lean mass is available, 47 sites associated with lean mass in the mothers in ALSPAC at p<1e-04 are available in the EWAS catalog (36); http://www.ewascatalog.org/. We checked for inflation of these sites in our data using QQ plots and lambda values, and tested enrichment for these sites using a Wilcoxon rank sum test.

## Results

### Study characteristics

Of the 1869 individuals with EPIC array DNAm profiles passing QC, we removed 267 because they were either of non-white ethnicity or had missing ethnicity data – this was because non-white ethnicity was associated with lower rates of smoking for both maternal and paternal grandmothers (p=0.03 and 0.007, respectively). Of the remaining 1602 participants, 285 were removed because their mother reported that she smoked during her pregnancy, and 73 further individuals were removed because they reported smoking themselves. Of the 910 individuals with 450k DNAm data passing QC and filtering, 125 individuals were removed because their mother reported smoking during pregnancy, and a further 59 were removed as they reported smoking themselves. All individuals in the 450k dataset were of white ethnicity. Table 1 summarises the characteristics of the adolescent datasets; supplementary table 1 details the numbers of participants with complete data in each EWAS.

### Discovery EWAS results

No associations tested using the main model (all covariates) survived the Bonferroni-adjusted p-value threshold (p<9e-8). variation. All associations p<1e-04 using the main model are reported in supplementary tables 2-7.

### Replication

All associations in the replication dataset p<1e-04 using the main model are reported in supplementary tables 8-13. Firstly we tested replication at p<0.05/25 for each of the six EWAS. For maternal grandmother smoking, the association at a single site on the X chromosome replicates in the females only analysis (cg19782749, p=0.001; **Error! Reference source not found.**). For paternal grandmother smoking, none of the associations at the top 25 sites replicate. Secondly, we evaluated correlation of effect sizes for associations at the top sites in each EWAS. For maternal grandmother smoking in the all-individuals analysis, there were moderate negative correlations for the top 25 to 200 sites (R=-0.18 to −0.45, p<0.03). Within each of the other five EWAS analyses, moderate correlations (R=0.28 to 0.45, p<0.05) were found for at most two two subsets of sites - correlations were otherwise small (R<0.2), and four analyses featured both negative and positive correlations. Therefore we do not find consistent evidence of replication of effect sizes in our analyses. Thirdly, we asked if direction of effect was preserved in replication data for the top associated sites. There was again evidence supporting replication for maternal grandmother smoking in female grandchildren (in four of five tests p<0.009, binomial test), but none of the other EWAS. Details can be found in supplementary table 14.

Using the same replication methods we evaluated agreement between associations observed in male and female stratified analyses in the discovery dataset. For maternal grandmother smoking, effects of associations at top female sites appear to be negatively correlated with effects in males (R = −0.21 to −0.43, p < 0.05). None of the other replication analyses yielded evidence for agreement or disagreement between top male and female associations. For paternal grandmother smoking, effect sizes of top female sites were positively correlated with effects in males (R = 0.34 to 0.66, p < 0.04); we also observe agreement in direction of effect for associations at the top 200 female sites. Details are in supplementary table 15.

### Meta-analysis

As the datasets were generated using different Illumina Beadchip arrays, we meta-analysed only the 438,459 sites that were common to both arrays. No associations survived Bonferroni-adjustment for multiple tests (p < 2.4e-07).

### X chromosome

When testing the X chromosome, only one association survived adjustment for multiple tests (p < 2.7e-6). The association was with paternal grandmother smoking in the males (cg27456137; p=1.9e-06); **Error! Reference source not found.**. The probe for this site has been flagged (23) as cross-hybridising to a 49bp sequence 500bp from cg27456137. Three probes on the EPIC array reside within that 49bp sequence; however none were associated with either grandmother smoking near genome-wide significance (all p>0.03).

### Escapees

When testing whether DNAm sites located within escapee regions were associated at the Bonferroni corrected p-value p<1.5e-06 in the discovery dataset, we find no sites associated with maternal or paternal grandmother smoking.

### Imprinting control regions

We similarly tested the hypothesis that transmission might involve sites within ICRs. We observe one association that survives correction for multiple tests (p<5.1e-05); the association is with paternal grandmother smoking (cg15068552, p=2.2e-05); **Error! Reference source not found.**.

### Testing associations and replication earlier in life

In cord blood, we find one site associated with maternal grandmother smoking in all individuals, and two sites associated with paternal grandmother smoking in females (see **Error! Reference source not found.** for a summary). In the 7-year-olds, no sites were associated with either grandmother smoking in any of the six analyses. All associations p>1e-04 using the main model are reported in supplementary tables 16-27. None of these associations were observed at adolescence (i.e., in the main discovery dataset) below the p<0.05/3 threshold (all p>0.07).

We then tested whether two of the three associations observed at adolescence (i.e., in the main discovery dataset) were observed at birth and at 7 years (cg15068552 in all individuals when the paternal grandmother smoked, and cg19782749 in females when the maternal grandmother smoked; cg27456137 could not be tested because it was not measured by the 450k array). We see a suggestion of replication at cg15068552 at birth in all individuals when the paternal grandmother smoked (p=0.02), and at cg19782749 at 7 years in females when the maternal grandmother smoked (p=0.04).

### Transcription factor binding site analysis

Using locus overlap enrichment analysis (LOLA), we find enrichment of the top 25 EWAS associations at the TFBS for four TFs (nominal p < 0.05) in EWAS of paternal grandmother smoking. CtBP2 is enriched for the EWAS of males and females (log OR=1.8, p=0.02); NR2F2 is enriched in the female-stratified EWAS (log OR=1.8, p=0.02); and CtBP2 (log OR=1.9, p=0.01), SAP30 (log OR=1.4, p=0.04), and ZKSCAN1 (log OR=2.3, p=0.02) are enriched in the males-only EWAS. There are no enrichments in the maternal grandmother smoking analyses. These enrichment results are illustrated in **Error! Reference source not found.**. We then used the UCSC genome browser to assess whether the six individual sites identified were located within TFBS. We find all six are located in sites which bind at least one TF; these are detailed in supplementary table 28.

### Enrichment of prenatal- and own smoking- associated sites

Among sites associated with prenatal smoking, we observe some inflation for associations with paternal grandmother smoking in males (lambda=1.27 ± 0.13) and females (lambda=1.12 ± 0.11). This inflation is replicated only for males in the 450k dataset (lambda=1.46 ± 0.12). Among sites associated with own smoking, there is weak inflation for associations with paternal grandmother smoking in females (lambda=1.16±0.05), but this association is not replicated. Inflation results are summarised in Table 3.

**Table 3:**
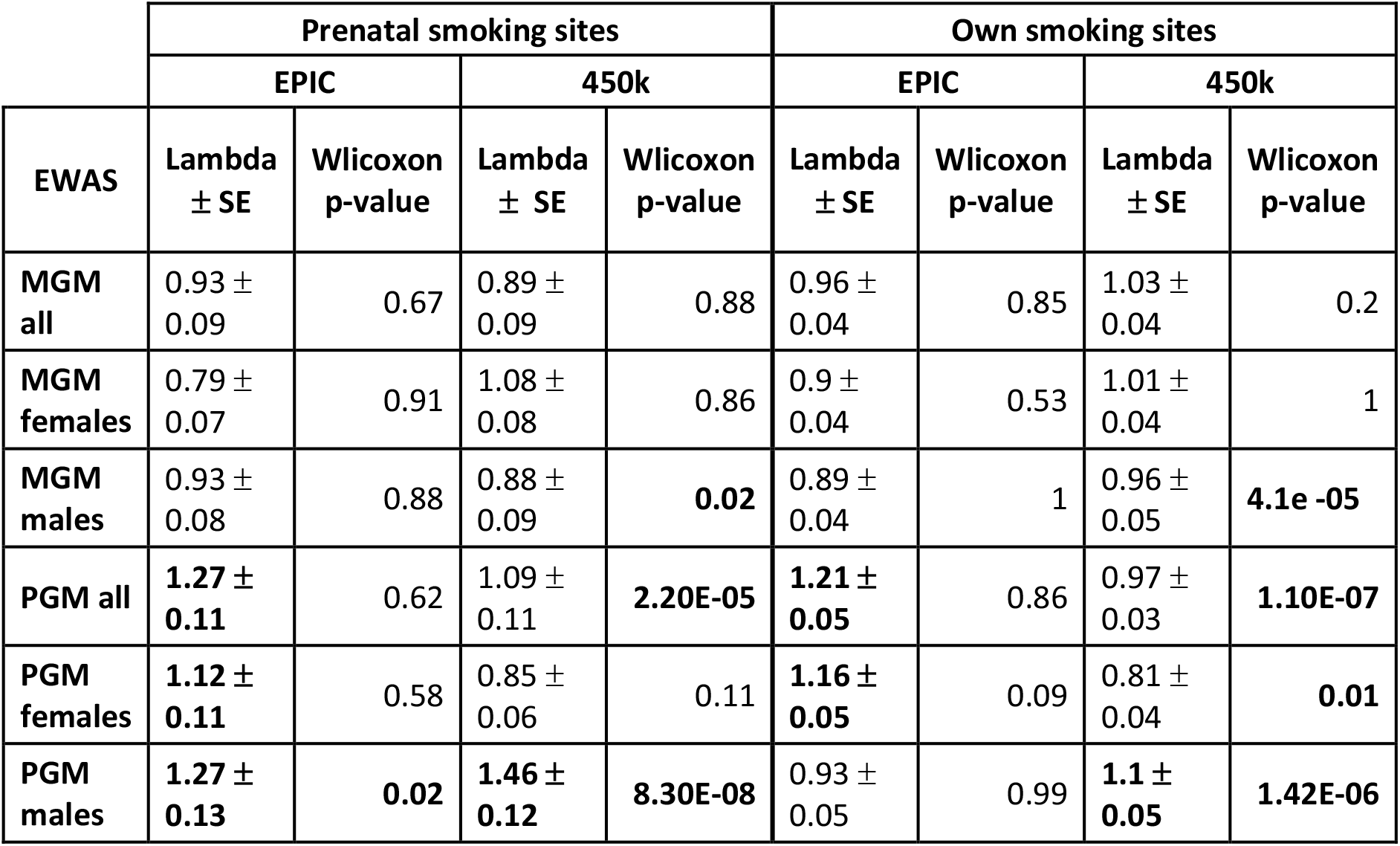
Summary of lambda values to illustrate inflation, and Wilcoxon rank-sum p-values to illustrate enrichment, of DNAm sites previously identified to be associated with prenatal smoke exposure and own smoking.

| EWAS | Prenatal smoking sites |  |  |  | Own smoking sites |  |  |  |
| --- | --- | --- | --- | --- | --- | --- | --- | --- |
|  | EPIC |  | 450k |  | EPIC |  | 450k |  |
|  | Lambda<br>± SE | Wlcoxon<br>p-value | Lambda<br>± SE | Wlcoxon<br>p-value | Lambda<br>± SE | Wlcoxon<br>p-value | Lambda<br>± SE | Wlcoxon<br>p-value |
| **MGM**<br>**all** | 0.93 ±<br>0.09 | 0.67 | 0.89 ±<br>0.09 | 0.88 | 0.96 ±<br>0.04 | 0.85 | 1.03 ±<br>0.04 | 0.2 |
| **MGM**<br>**females** | 0.79 ±<br>0.07 | 0.91 | 1.08 ±<br>0.08 | 0.86 | 0.9 ±<br>0.04 | 0.53 | 1.01 ±<br>0.04 | 1 |
| **MGM**<br>**males** | 0.93 ±<br>0.08 | 0.88 | 0.88 ±<br>0.09 | **0.02** | 0.89 ±<br>0.04 | 1 | 0.96 ±<br>0.05 | **4.1e -05** |
| **PGM all** | **1.27 ±**<br>**0.11** | 0.62 | 1.09 ±<br>0.11 | **2.20E-05** | **1.21 ±**<br>**0.05** | 0.86 | 0.97 ±<br>0.03 | **1.10E-07** |
| **PGM**<br>**females** | **1.12 ±**<br>**0.11** | 0.58 | 0.85 ±<br>0.06 | 0.11 | **1.16 ±**<br>**0.05** | 0.09 | 0.81 ±<br>0.04 | **0.01** |
| **PGM**<br>**males** | **1.27 ±**<br>**0.13** | **0.02** | **1.46 ±**<br>**0.12** | **8.30E-08** | 0.93 ±<br>0.05 | 0.99 | **1.1 ±**<br>**0.05** | **1.42E-06** |

### Inflation test for lean mass-associated DNAm sites

We observe no evidence for inflation among CpG sites associated with lean mass. These inflation results are summarised in supplementary table 29.

## Discussion

In summary, we find some evidence for effects of grandmother smoking on DNA methylation in her adolescent grandchildren; on the X chromosome, in an ICR, in TFBS, and among prenatal smoking-associated DNAm sites. We also find three sites associated with grandmaternal smoking in cord blood, but these associations do not appear to persist. In most cases, associations appear to be sex-specific in line with previous research (8–10). Associations are summarised in Figure 2.

**Figure 1:**
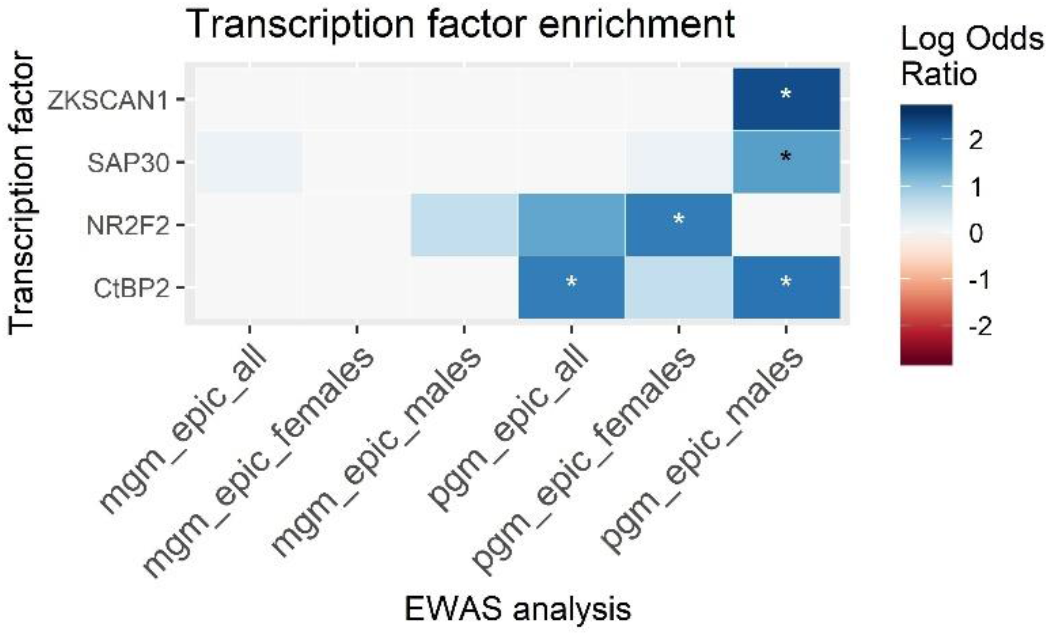
Transcription factor binding site enrichments for TFBS that reached p<0.05 for at least one of the EWAS. Heatmap is coloured by the log odds ratio, *=p<0.05

**Figure 2:**
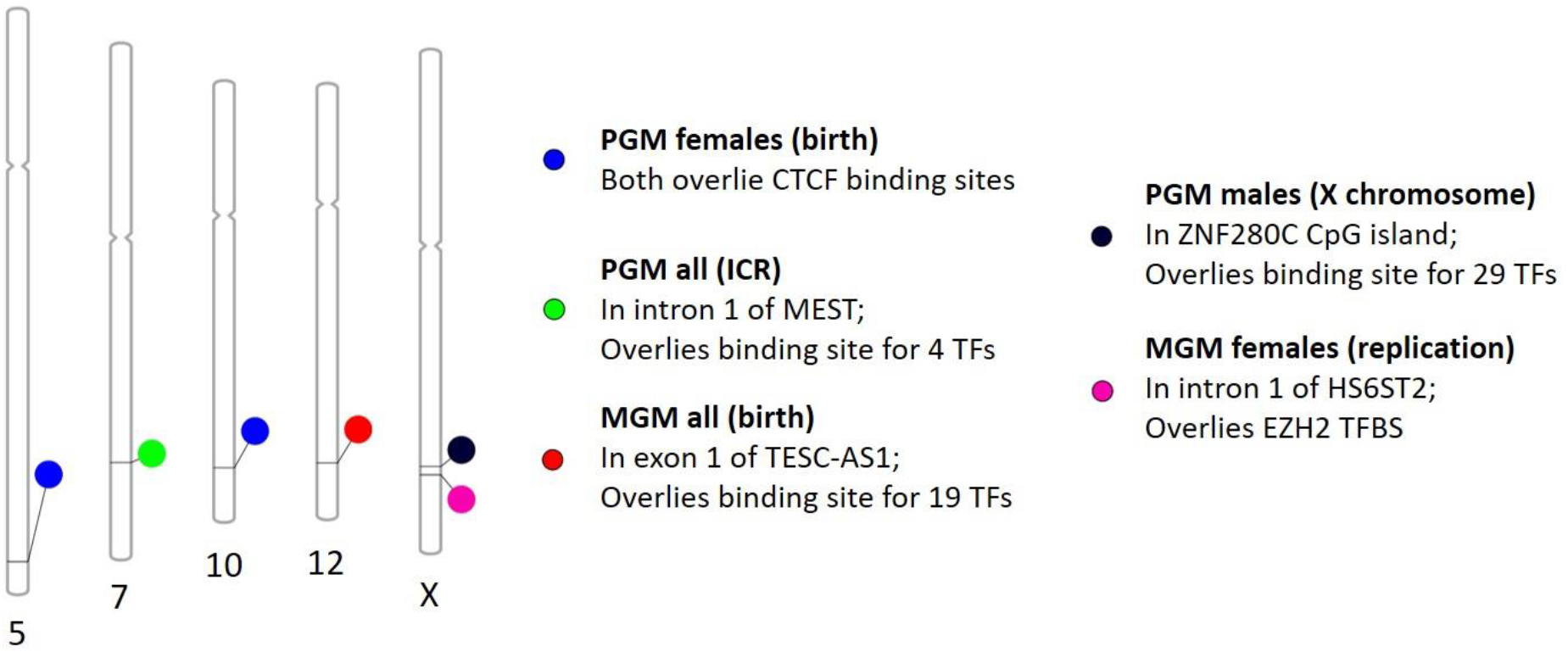
Graphical summary illustrating the main study findings. MGM = maternal grandmother, PGM = paternal grandmother. Bold text indicates EWAS (analysis); text indicates whether the site is located within a gene and whether it is located in a TFBS.

We find some evidence for mechanisms by which DNAm might be preserved through phases of de- and re-methylation in the germ cell. Two of the six sites we identify are on the X chromosome, giving a possible route by which sex-specific differences in transmission of responses across generations might occur. We find evidence suggesting TFs might have a role in the transmission of epigenetic responses to smoking across generations - both from the enrichment analysis, and the location of all six individual sites within TFBS. We find evidence for a single site residing within an ICR, but find no evidence for sites in regions known to escape de-methylation in germ cells. Finally, we find suggestive evidence of replication of two sites identified in adolescents in earlier DNAm samples (one at birth and one at 7 years), although no site replicates across all three timepoints.

We find evidence of inflation and enrichment of sites associated with prenatal smoking only in males when their paternal grandmother smoked, and do not find consistent inflation of sites associated with own smoking. This could suggest that grandmaternal smoking affects DNAm through different mechanisms to maternal smoking. The inflation we see in males is contrary to previous null prenatal findings (18); the reason for this discrepancy may be that we test a larger number of sites. We do not see any inflation or enrichment of lean mass associated DNAm sites in our analyses, suggesting that the differences in lean mass observed previously (13) may not be related to differences in DNAm.

Because TFBS are a consistent feature of our findings, our study supports the idea that DNAm changes may be linked to ancestral smoking by TF binding events. These binding events could either shield DNAm from being modified in early development or induce DNAm changes consistent with ancestral smoking, as DNAm status can be restored by TFs during germline and embryonic development following erasure (6, 7). However it is not clear why the associations we do see would change over time, and so we cannot rule out the possibility that we find differences at these DNAm sites due to another factor that is influenced by grandmaternal smoking, such as parental behaviour. We suggest TFBS might present the most promising line of future work in transgenerational epigenetic responses in humans.

Strengths of our study are that we assessed grandmaternal smoking effects in a large cohort of humans with ancestral smoking data, alongside rich phenotypic data. We have DNAm data from birth so were able to assess whether DNAm differences at these sites are present between birth and adolescence. Limitations include that the 450k and EPIC array platforms only cover around 2% and 4% of the genome, respectively, and that our replication dataset came from the same birth cohort as the discovery data.

## Acknowledgements

We are extremely grateful to all the families who took part in this study, the midwives for their help in recruiting them, and the whole ALSPAC team, which includes interviewers, computer and laboratory technicians, clerical workers, research scientists, volunteers, managers, receptionists and nurses.

