## Supplementary methods for "Grandmaternal smoking during pregnancy is associated with differential DNA methylation in their grandchildren"

#### Affiliations

### Supplementary methods

#### Cohort description

ALSPAC is a pre-birth cohort designed to determine the environmental and genetic factors that are associated with health and development of the study offspring (1-3). Ethical approval for the study was obtained from the ALSPAC Ethics and Law Committee and the Local Research Ethics Committees (Birmingham, 2018). Consent for biological samples has been collected in accordance with the Human Tissue Act (2004). Informed consent for the use of data collected via questionnaires and clinics was obtained from participants following the recommendations of the ALSPAC Ethics and Law Committee at the time.

Pregnant women resident in Avon, UK with expected dates of delivery 1st April 1991 to 31st December 1992 were invited to take part in the study. The initial number of pregnancies enrolled is 14,541 (for these at least one questionnaire has been returned or a "Children in Focus" clinic had been attended by 19/07/99). Of these initial pregnancies, there was a total of 14,676 fetuses, resulting in 14,062 live births and 13,988 children who were alive at 1 year of age.

When the oldest children were approximately 7 years of age, an attempt was made to bolster the initial sample with eligible cases who had failed to join the study originally. As a result, when considering variables collected from the age of seven onwards (and potentially abstracted from obstetric notes) there are data available for more than the 14,541 pregnancies mentioned above.

The number of new pregnancies not in the initial sample (known as Phase I enrolment) that are currently represented on the built files and reflecting enrolment status at the age of 24 is 913 (456, 262 and 195 recruited during Phases II, III and IV respectively), resulting in an additional 913 children being enrolled. The phases of enrolment are described in more detail in the cohort profile paper and its update (2, 3). The total sample size for analyses using any data collected after the age of seven is therefore 15,454 pregnancies, resulting in 15,589 fetuses. Of these 14,901 were alive at 1 year of age.

Please note that the study website contains details of all the data that is available through a fully searchable data dictionary and variable search tool: http://www.bristol.ac.uk/alspac/researchers/our-data/

ALSPAC fully supports Wellcome and the RCUK policies on open access. The process for obtaining access to data is described on the study website: http://www.bristol.ac.uk/alspac/researchers/data-access/.

#### Study exclusions

All adolescents whose mother smoked during pregnancy, and all adolescents who reported smoking themselves, were removed from our analyses to ensure as far as possible that only effects arising from the grandmother smoking would be detected. Maternal smoking during pregnancy was assessed using a previously defined measure (4), which identified whether the mother had smoked in any of the three trimesters of that pregnancy. Own smoking was also represented by a composite measure (5); the adolescents were classified as smokers if they reported smoking more than one cigarette per week at the 15 year clinic and classified as non-smokers if they reported that they had never tried a cigarette.

As a small number of the participants with EPIC array data were of non-white ethnicity (ALSPAC reports ethnicity as either white or non-white, derived from parental self-report), we conducted a sensitivity analysis to ascertain whether we could include individuals of all ethnicities in our analyses.

Ethnicity is a social construct so we would not expect transmission of biological responses across generations to differ across ethnicities; however as reduced rates of both maternal and paternal grandmother smoking were associated with non-white ethnicity (p=0.03 and 0.007, respectively), we removed individuals of non-white ethnicity from the analysis.

#### Ancestral smoking data

The mothers and fathers of the index children of ALSPAC each answered a questionnaire about their parents (the grandparents of the ALSPAC index children). The questionnaire included whether their mother (the study child's maternal or paternal grandmother) smoked. If the study parent reported their mother did smoke, they were asked a further question about whether she smoked whilst she was pregnant with them (the study parent). Possible responses to this further question were yes/no/don't know, where 'don't know' meant the study grandmother smoked but the study parent was unsure whether she smoked whilst pregnant with them. Consistent with our previous work (6-8), we have conducted our analysis with the assumption that the grandmothers in the 'don't know' category did smoke whilst they were pregnant with the study mother or father. This assumption was validated by the demonstration that the study parents in the 'don't know' category had reduced mean birthweight in comparison to those whose mother had definitely not smoked during pregnancy (8). Therefore, responses to smoking for each grandmother were coded as a categorical 'Yes' or 'No' in our study. For all analyses we kept maternal and paternal grandmothers separate.

#### Assessing DNAm sites for genetic and trait associations

To assess the sites we identified in our analyses, we checked whether they had been associated with genetic variants and with traits in previous studies. To test for association with genetic variants, we used the GoDMC (9) meta-analysis database (http://mqtldb.godmc.org.uk/), which contains all *cis* and *trans* CpG-SNP associations from the largest methylation quantitative trait loci (mQTL) study to date. To test for association with traits in previously published literature, we used the EWAS catalog (10); http://www.ewascatalog.org/.

#### Summarising associations

A summary plot was produced using the PhenoGram (11) web resource.

### Supplementary results

#### Associations between covariates and grandmaternal smoking

The only notable associations between covariates and grandmaternal smoking in the adolescent data is with predicted eosinophil cell type proportions; in the discovery dataset, proportions are higher in females when the paternal grandmother smoked (p=0.004). In the replication dataset, eosinophil proportions are lower in all individuals (p=0.03) and in males (p=0.02) when the paternal grandmother smoked, and lower in females when the maternal grandmother smoked (p=0.04). At birth, lower gestational age was associated with paternal grandmother smoking in all individuals (p=0.004) and females (p=0.0007). In the 7-year-olds, decreased CD4T and increased monocyte cell proportions were associated with paternal grandmother smoking (p=0.03 and 0.05, respectively).

#### Assessing DNAm sites for genetic and trait associations

We find that three of the six sites associated with grandmaternal smoking are mQTLs – one is a *cis* mQTL and two are *trans* mQTLs. One of the sites associated with grandmaternal smoking in cord blood (cg22682200) has previously been associated with gestational age, and in that specific analysis in our study gestational age was associated with paternal grandmaternal smoking status in females at birth. This may suggest residual confounding of gestational age. The other site associated with paternal grandmother smoking in cord blood in females has previously been associated with sex. There are no associations between the DNAm sites and traits previously associated with grandmaternal smoking.

| **CpG** | **Chr** | **Analysis** | **SNP associations** | **Trait associations** |
|---|---|---|---|---|
| cg19426678 | 12 | Birth – MGM all | No associations | HIV infection (p=2.1e-06) |
| cg22682200 | 10 | Birth – PGM females | 57 *trans* associations | Tissue type (fetal vs adult liver (p=1.1e-22); blood vs buccal (p=1.9e-10)); gestational age (p=3.5e-10) |

| | | | | |
|---|---|---|---|---|
| cg26827966 | 5 | Birth – PGM females | No associations | blood vs buccal (p=6.2e-34); sex (p=4.7e-08 and 1.5e-07) |
| cg19782749 | X | Adolescence – MGM females replication | 213 *cis* associations | Sex (4.1e-202) |
| cg27456137 | X | Adolescence – PGM males Xchr | No associations | Papuan ancestry (p=3.4e-5) |
| cg15068552 | 7 | Adolescence – PGM all ICR | 205 *trans* associations | Clear cell renal carcinoma (p=8.6e-33) |

*Table 1: SNP and trait associations for the six DNAm sites associated with grandmaternal smoking*
